# Lungfish comparative genomics reveals ancient gene networks co-opted for life on land

**DOI:** 10.64898/2026.03.20.713117

**Authors:** Klara Eleftheriadi, Judit Salces-Ortiz, Nuria Escudero, Carlos Vargas-Chávez, Ryan D. Heimroth, Laia Ribas, Mercedes Blázquez, Salvador Carranza, Irene Salinas, Rosa Fernández

**Affiliations:** Institute of Evolutionary Biology (CSIC-UPF), Barcelona (Spain); Center for Evolutionary and Theoretical Immunology, Department of Biology, University of New Mexico, Albuquerque, New Mexico (USA); Institut de Ciències del Mar, Consejo Superior de Investigaciones Científicas (ICM-CSIC), Barcelona (Spain)

## Abstract

The transition from water to land was a defining event in vertebrate evolution, requiring major changes in physiology and gene regulation. Lungfish, the closest extant relatives of tetrapods, offer a unique window into this process through their capacity for estivation, a physiological state that mirrors key demands of terrestrial life. Here, we constructed and compared gene co-expression networks across five vertebrate species using multi-tissue transcriptomic data. Focusing on estivating lungfish tissues, we identified conserved modules enriched in core functions, including metabolic regulation, RNA processing, and protein turnover. We further show that a subset of retained ohnologs from the vertebrate 2R-WGD act as network hubs under directional selection in key phylogenetic branches. These genes, involved in membrane trafficking, cytoskeletal regulation, water balance, and stress response, illustrate how duplicated, dosage-sensitive genes were co-opted to meet the physiological demands of terrestrial adaptation. Together, our findings support a model in which ancient whole-genome duplications supplied raw material later shaped by selection and network rewiring to enable extreme physiological adaptations such as estivation.

**Teaser:** Ancient duplicated gene networks in lungfish estivation reveal how vertebrates repurposed genomic toolkits to adapt to land.

## Introduction

Vertebrate terrestrialization, the transition from an aquatic to terrestrial environment, marks one of the most pivotal chapters in the history of life on Earth. Lungfish (Dipnoi) are lobe-finned fish (Sarcopterygii) with a fossil record dating back to the Early Devonian period (*1*). Lungfish are the closest extant relatives to tetrapods (*2*). Although they were abundant during the Devonian period, only six species survive today, making them true “living fossils” (*3*). As obligate air-breathers capable of surviving prolonged droughts through estivation, lungfish possess physiological adaptations that parallel the key challenges faced during terrestrialization, including dehydration tolerance, coping with oxygen variability, and metabolic suppression (*3*, *4*). Estivation may thus represent an exaptive trait, offering a functional window into the preconditions for terrestrial life. While the fossil record of extinct transitional taxa like *Tiktaalik roseae* have clarified the morphological progression from lobe-finned fishes to early tetrapods (*5*, *6*), extant taxa such as lungfish are essential for probing the physiological and genetic mechanisms underlying this transition.

Vertebrates underwent two rounds of whole genome duplication (2R-WGD) early in their evolution, dramatically expanding the genomic repertoire (*7*, *8*). Although most duplicated genes have been lost, retained ohnologs are enriched for dosage-sensitive, regulatory, and developmental functions (*9*, *10*). These retained duplicates often undergo sub- or neo-functionalization and are hypothesized to play a role in lineage-specific adaptations (*11*, *12*). These duplications likely expanded the genomic toolkit available to early vertebrates, but whether retained ohnologs directly contributed to terrestrialization remains elusive. Their dosage-sensitive and regulatory roles make them key potential candidates, yet their specific co-option into adaptations such as water balance, stress response, and metabolic regulation has not been systematically tested.

Comparative genomic and transcriptomic studies have positioned lungfish as a pivotal lineage for understanding vertebrate terrestrialization. The genome of the Australian lungfish (*Neoceratodus forsteri*) revealed extensive synteny with tetrapods, expansions in gene families linked to air-breathing and olfaction, and conservation of limb-development regulators (*13*). More recently, the genomes of the African (*Protopterus annectens*) and the South American (*Lepidosiren paradoxa*) lungfish have shed light on transposon-driven genome expansion and chromosomal conservation with tetrapods (*14*). Transcriptomic resources (*15*, *16*) and single-cell atlases of respiratory organs (*17*) have enabled cell-type and pathway comparisons across sarcopterygians, while proteomic analyses have examined terrestrialization-linked changes in skin mucus (*18*). These studies collectively underscore lungfish as a model for bridging the gap between aquatic and terrestrial vertebrates.

Despite these advances, no study has yet integrated gene expression across multiple tissues and physiological systems, particularly from estivating lungfish, into a comparative framework spanning aquatic and terrestrial vertebrates. This is critical because African lungfish undergo whole-body, profound structural, metabolic and physiological adaptations during water-land transitions. For instance, previous work uncovered transcriptional and histological adaptations in diverse organs such as the skin, brain, heart, kidney, gut and lungs (*19–24*). As such, critical questions remain unanswered: Do estivation-associated genes represent ancient, conserved toolkits or more recent lineage-specific innovations? Are there shared transcriptional programs across tissues and organs that represent a pan-body response to land adaptation? Have ohnologs retained from 2R-WGD been systematically co-opted into the physiological strategies needed for terrestrial life? And to what extent has directional selection sculpted these duplication-enriched networks during key evolutionary transitions?

Here, we address these questions by performing a multi-species, multi-tissue gene co-expression network analysis to investigate the molecular underpinnings of vertebrate terrestrialization. Using the African lungfish as a focal taxon, we integrate gene expression of multiple tissues (gills, lungs, kidney, brain, intestine, muscle, skin, heart, gut and liver) from water and land-adapted states including comparative data from two teleosts (the zebrafish (*Danio rerio*) and the European sea bass (*Dicentrarchus labrax*), an amphibian (the natterjack toad, *Epidalea calamita*), and a reptile (the moorish gecko, *Tarentola mauritanica*). These species were selected to represent key extant lineages spanning the water-to-land transition - fully aquatic (freshwater and marine teleosts), amphibious (amphibians) and fully terrestrial (reptiles), providing a phylogenetic framework to distinguish ancestral traits from those driven by terrestrial adaptation. This framework allowed us to test whether conserved modules, retained ohnologs, and directionally selected genes intersect in ways that could have facilitated terrestrial adaptation. Specifically, we inferred species-specific gene co-expression modules via Weighted Gene Co-expression Network Analysis (WGCNA), identified highly connected genes within the modules (hubs) and investigated the evolutionary origin of hub-containing hierarchical orthologous groups (HOGs) through phylostratigraphy. We then evaluated whether these lungfish hubs, particularly those expressed during estivation, are enriched for retained ohnologs from the 2R-WGD and show signatures of directional selection. Our novel, systems-level approach allows to pinpoint hub genes as central players governing the coordinated physiology of estivation and to highlight their potential role as genomic substrates for vertebrate terrestrial adaptation.

## Results & Discussion

### Conserved co-expression modules across vertebrate tissues

Understanding the evolution of complex traits, such as the vertebrate transition from aquatic to terrestrial life, requires moving beyond gene-by-gene comparisons and instead examining how groups of genes function together as coordinated regulatory units. Gene co-expression networks, inferred here by WGCNA, provide a systems-level framework to capture such coordinated gene behavior (*25*, *26*). These networks identify gene modules with similar expression, often corresponding to biological pathways or shared transcriptional regulation, making them ideal for studying traits driven by multi-gene programs (*27*).

To assess the evolutionary conservation of gene modules involved in terrestrial adaptation, we first sampled multiple tissues (gills, lungs, kidney, brain, intestine, muscle, skin, heart, gut, liver) across five key vertebrate species: two ray-finned fishes (the European sea bass and the zebrafish), the African lobe-finned lungfish, an amphibian (the natterjack toad), and a reptile (the moorish gecko) (Table 1). Lungfish tissue samples were obtained from free-swimming individuals as well as from laboratory-estivated fish (see Methods). The transcriptomes of the different tissues were sequenced using short-read Illumina RNA-seq (Tables S2-S6). In parallel, we generated long-read Iso-Seq data from pooled RNA across all tissues per species (Table S7) to generate *de novo* transcriptome assemblies, which were subsequently used as references for pseudomapping tissue-specific expression profiles (Figs S1-S5, Tables S2-S6). We then constructed species-specific gene co-expression networks using WGCNA, following scale free topology (Figs. S6-S10, Tables S8-S12) (Fig. 1), resulting in 37-63 modules per species (Figs. S11-S15). For each species we identified statistically significant modules per tissue (Figs. S11-S15, Tables S13-17). We focused our comparative analysis on modules derived from estivating tissues in lungfish, considering estivation a physiological proxy for preadaptation to terrestrial life, due to shared challenges such as dehydration, increased oxygen availability, and metabolic suppression (*18*). By integrating these datasets, we established a cross-species framework to compare co-expression architecture and to pinpoint modules that may underlie shared or lineage-specific physiological strategies. In particular, we reasoned that modules active during lungfish estivation would highlight candidate networks with relevance to terrestrial adaptation, since they capture regulatory responses to dehydration, metabolic suppression, and variable oxygen conditions, all hallmark challenges of life on land.

**Table 1.**
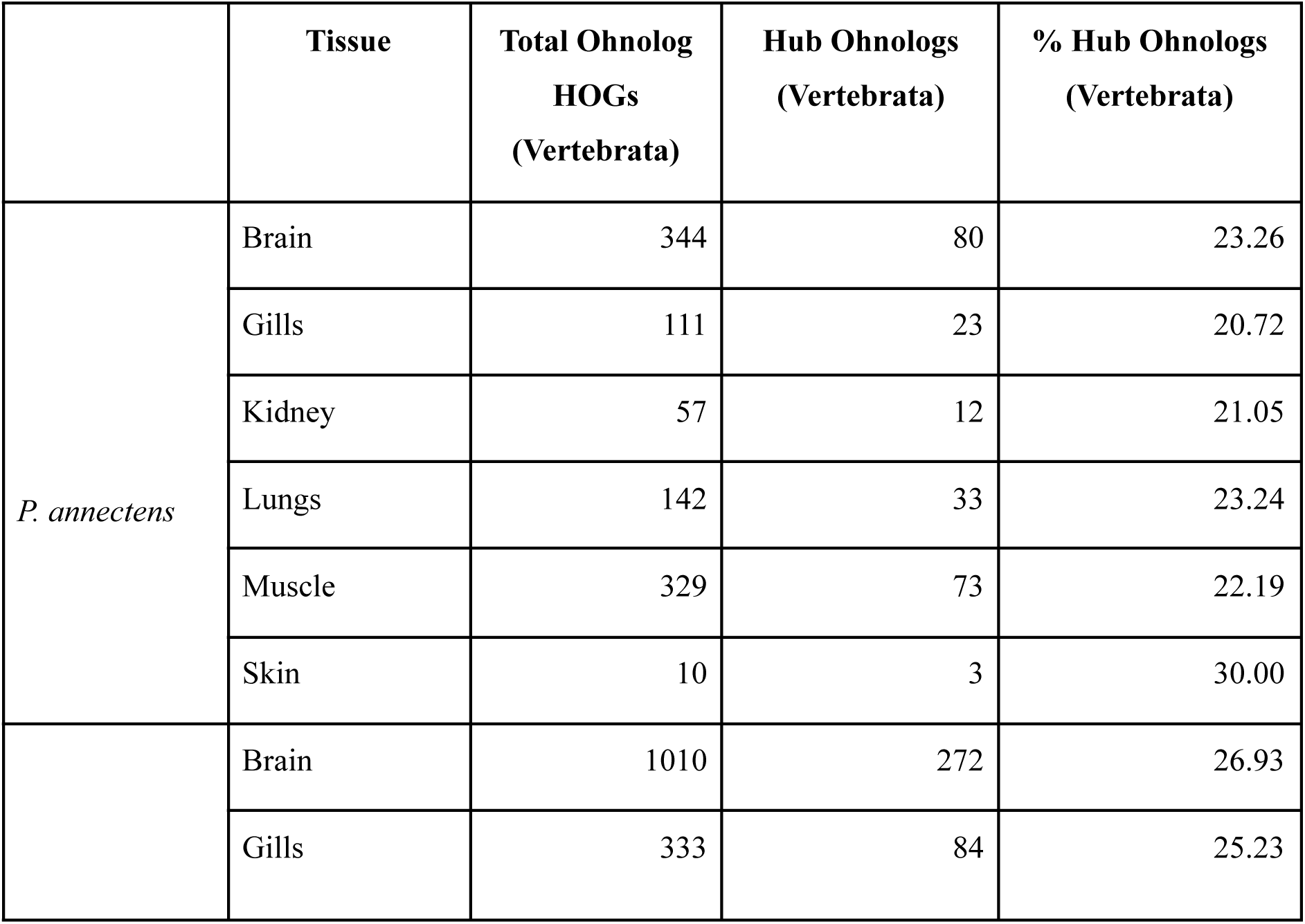
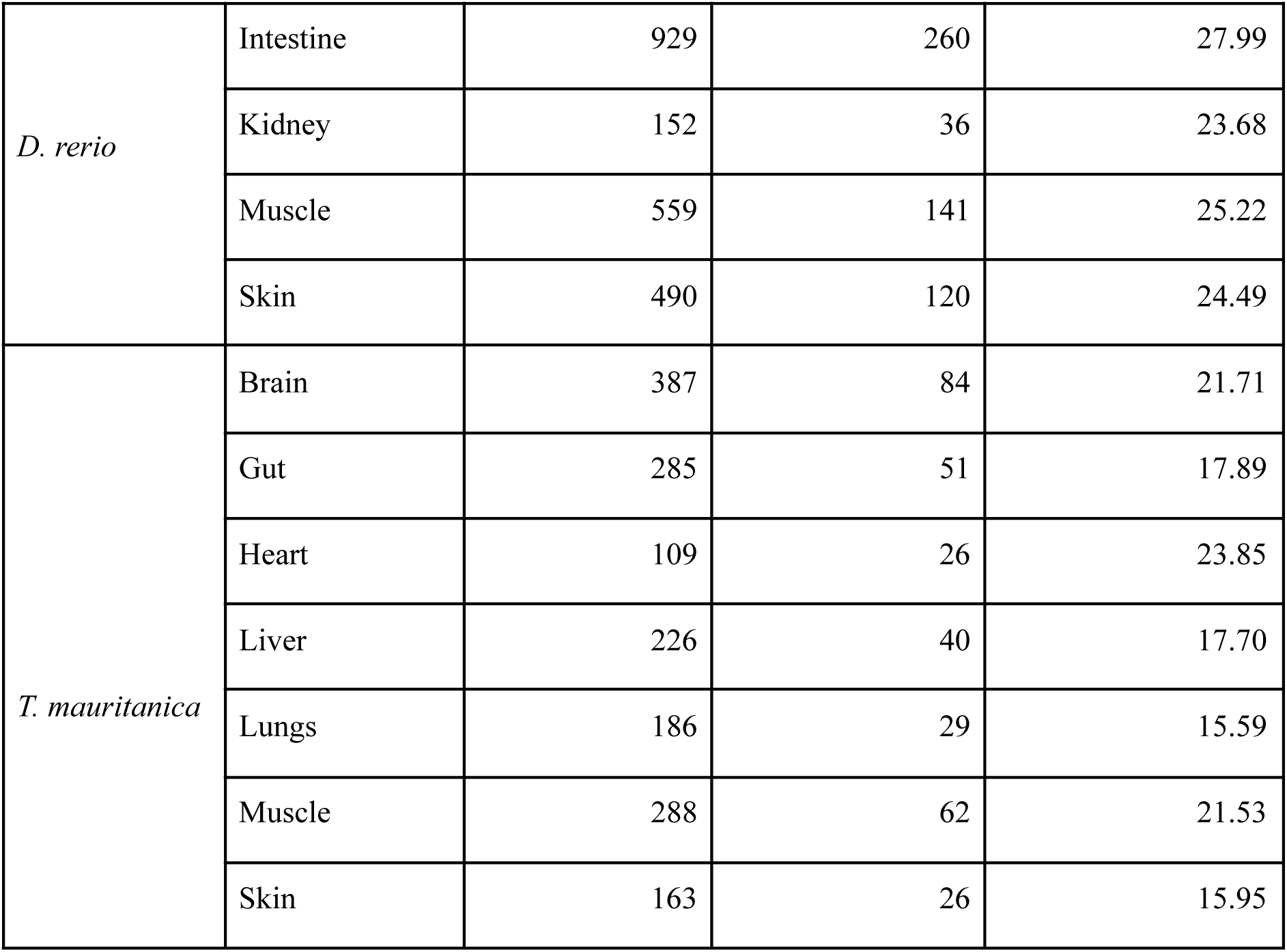
Retention of ohnologs in vertebrate-specific hub genes across tissues of lungfish, zebrafish, and the moorish gecko. The table shows, for each tissue, the total number of ohnolog-containing HOGs assigned to the Vertebrata phylostratum, the subset of those corresponding to hub genes, and the proportion of hub genes retained as ohnologs.

**Fig. 1.**
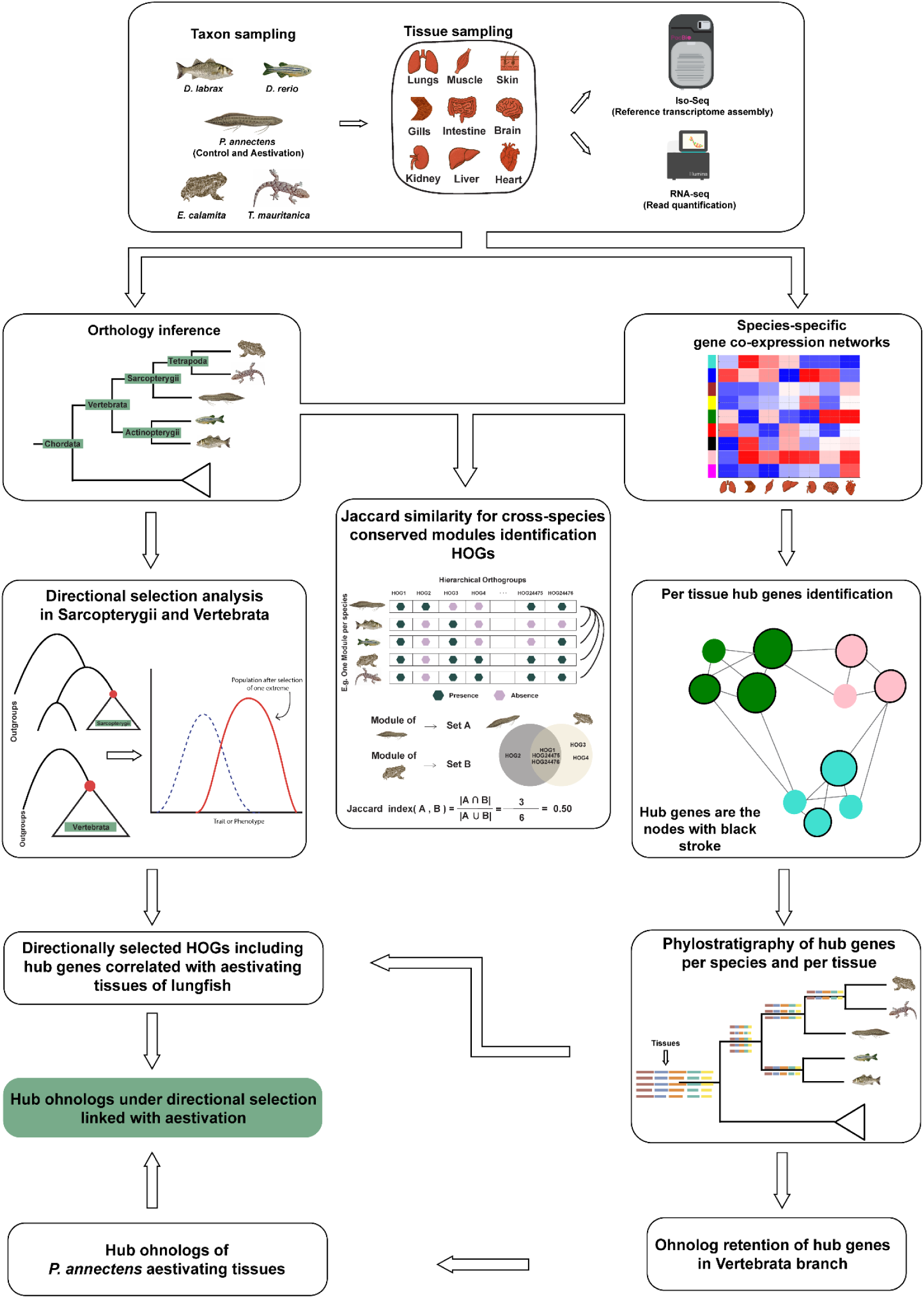
Schematic representation of the analytical workflow in this study. Overview of the species and tissues sampled, followed by analysis in three-axes: hub genes in co-expression networks, 2R-WGD ohnolog retention, and directional selection.Abbreviations: HOG, Hierarchical Orthologous Groups; 2R-WGD, two rounds of whole genome duplication.

To explore conservation patterns, we grouped genes from each species into Hierarchical Orthologous Groups (HOGs), using FastOMA (*28*), and then compared statistically significant lungfish estivation modules to those of other taxa by calculating pairwise Jaccard similarity scores based on shared orthologous genes (Fig. S19). This analysis revealed a few modules across species that show substantial compositional similarity (Jaccard index > 0.07) associated with brain, gill, kidney, and muscle tissues in lungfish (Fig. 2a, Fig. S13).

**Fig. 2.**
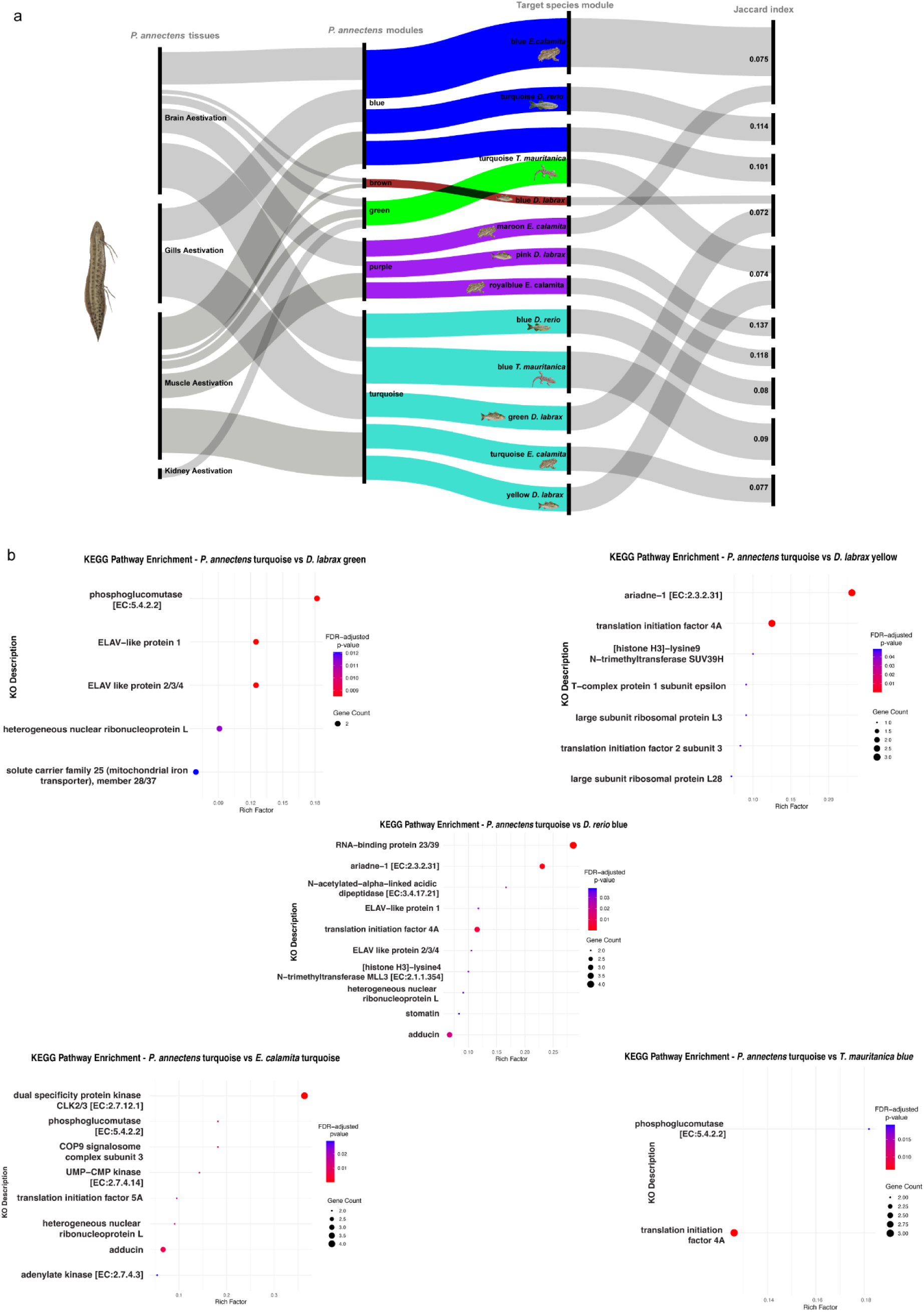
Statistically significant co-expressed modules for estivating tissues conserved across vertebrates. **(A)** Alluvial plot showing the lungfish modules under estivation with the highest Jaccard similarity to modules in other vertebrate species. The first column shows lungfish tissues in estivating animals with which the modules in the second column were significantly correlated. Second column, modules in lungfish estivating animals showing the highest degree of Jaccard similarity with other species, identified by colors. Third column, most similar modules between each species and estivating lungfish modules, identified by color. Fourth column, Jaccard index for each pairwise comparison. **(B)** KEGG pathway enrichment analyses of the lungfish turquoise module against the most similar modules in other species. Each dot represents an enriched KEGG pathway, with dot size indicating the number of genes, dot color the FDR-adjusted significance. The x-axis shows the enrichment ratio.

Of particular interest, the turquoise module of lungfish was the only one that showed convergent enrichment with modules from all other species, indicating a deeply conserved functional core (Fig. S19). This module was consistently enriched across taxa in KEGG pathways associated with metabolic regulation, translational control, RNA processing, protein turnover, cytoskeletal maintenance, and cellular transport (Fig. 2b). The genes driving these enrichments —including phosphoglucomutase and adenylate kinase (central to energy metabolism), ribosomal proteins and translation initiation factors (regulators of protein synthesis), hnRNP L and ELAV-like proteins (RNA splicing and mRNA stability), and the COP9 signalosome complex (protein turnover and protein ubiquitination control)— are well-established as fundamental for cellular function and adaptation to physiological stress (*29–34*). Notably, stomatin was also identified as enriched in this conserved module (Fig. 2b). While it is conserved between lungfish and zebrafish, recent evidence indicates it underwent relaxed selection in mammals transitioning back to the sea (*35*). Furthermore, it has also been suggested to have a relevant role in regulation of osmotic balance in erythrocytes, via encapsulation of aquaporin-1 and urea transporter-B in the erythrocyte membrane (*36*). This highly conserved module reflects the universal requirement of vertebrates for robust metabolic regulation, efficient gene expression, and cellular homeostasis, functions that underpin survival, physiological flexibility, and adaptation across diverse environments. The broad conservation of this module across both aquatic and terrestrial vertebrates, as well as its strong association with tissues integral to homeostasis (brain, gill, kidney, muscle) (Fig. 2a), suggests that these gene networks represent an ancestral genomic toolkit. Moreover, these findings indicate that such a toolkit likely predated the water-to-land transition, equipping early vertebrates with robust molecular mechanisms for physiological plasticity and stress tolerance. Thus, the turquoise module provides a compelling example of how deep conservation of gene networks can serve as a foundation for evolutionary innovations such as terrestrialization.

### Phylogenetic origin of network hub genes across vertebrates

We next assessed the evolutionary age of genes occupying central positions in these networks across vertebrates via phylostratigraphic analyses (Fig. 1). In brief, hub genes were assigned to HOGs, and their origins were mapped to each internode (phylostratum hereafter) to determine whether core regulatory components of these networks are evolutionarily ancient or represent lineage-specific innovations. Hub genes were first assigned to HOGs, and to avoid redundancy, we counted each HOG only once per tissue even if multiple hub genes from the same group were present. For each tissue and species, we calculated the proportion of hub-containing HOGs originating at each phylostratum, normalized by the total number of hub-containing HOGs in that tissue.

A large percentage (approximately 40 to 70%) of hub-containing HOGs traced back to deep ancestral nodes, especially the Metazoan/Deuterostome phylostratum (root), indicating that the regulatory architecture of these co-expression modules is evolutionarily ancient (Fig. 3, Table S18). Strikingly, we also observed a prominent enrichment of hub gene origin in the branch leading to Vertebrata, consistently across tissues and species (ca. 15 to 35%; Fig. 3). This pattern suggests that the expansion of regulatory complexity and network modularity likely began early in the vertebrate lineage, possibly as a result of gene duplications and regulatory innovations during the early diversification of vertebrates. Similar trends have been reported in plants and other lineages, where regulatory hubs and key innovations frequently trace back to genes that predate major evolutionary transitions, rather than arising solely from lineage-specific novelty (*37–40*).

**Fig. 3.**
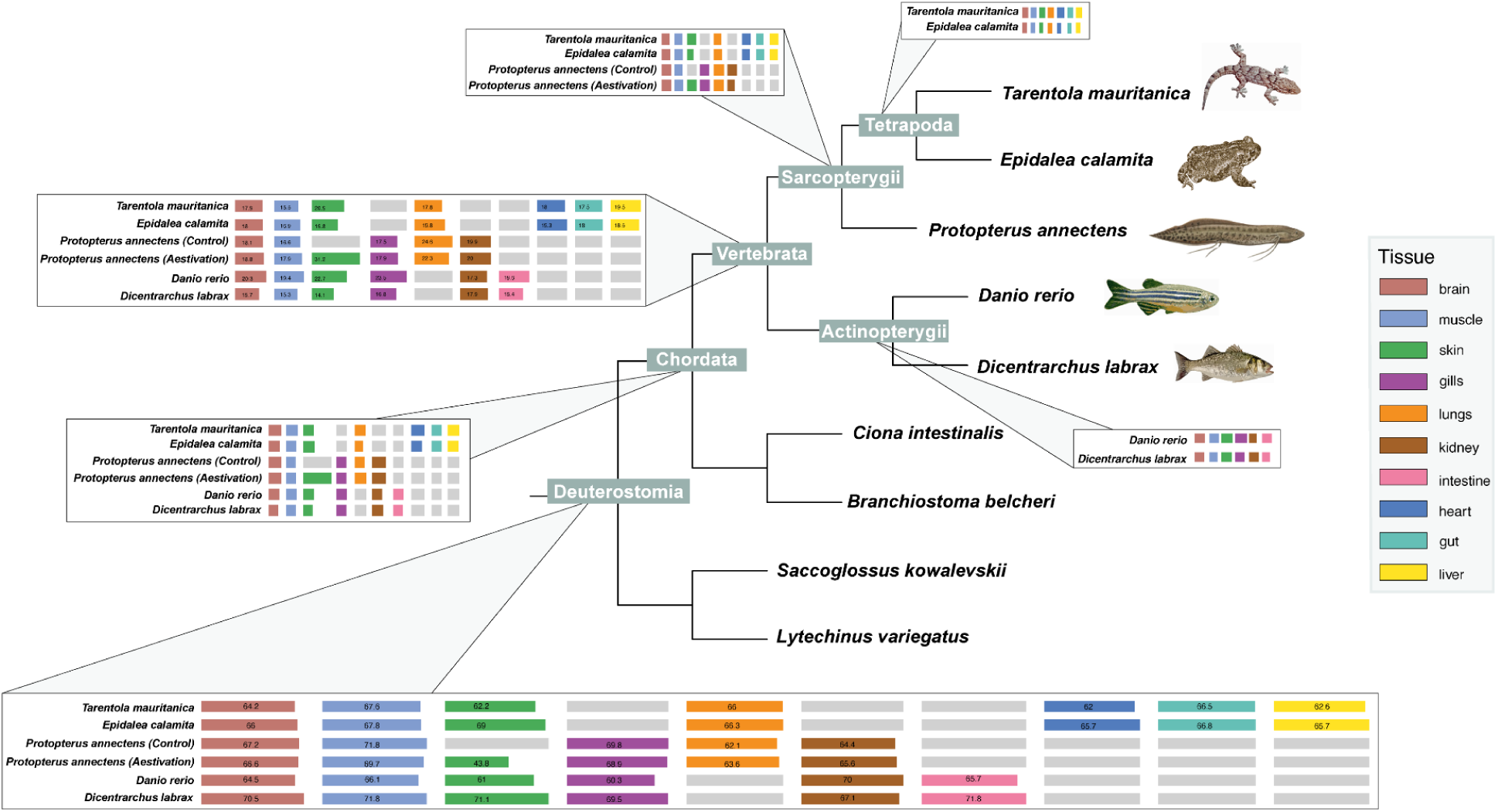
Evolutionary origin of hub genes across vertebrate tissues. Phylostratigraphic analysis showing the placement of hub genes per species in the corresponding phylostratum of origin based on HOGs. Colored boxes indicate the proportion of hub genes assigned to each phylostratum, color-coded by tissue. Two bursts of gained hub genes were observed, in the root of the tree and in the Vertebrata phylostratum.

### Ohnolog retention and directional selection in hub genes orchestrating estivation in lungfish

Whole-genome duplication (WGD) events have played a major role in shaping the genomic architecture of vertebrates by generating duplicated gene sets, ohnologs, that serve as a substrate for evolutionary innovation (*41*, *42*). While most gene duplicates are eventually lost, a subset is selectively retained, often due to their roles in complex regulatory networks or dosage-sensitive processes (*9*). Because the two rounds of whole-genome duplications (2R-WGD) are known to have shaped vertebrate genome complexity (*7*, *8*), we examined whether vertebrate-specific hub genes across species show elevated retention of ohnolog pairs.

We cross-referenced hub-containing HOGs assigned to the Vertebrata phylostratum with curated ohnologs from the OHNOLOGS v2 database (*10*) for three key lineages (zebrafish, lungfish and moorish gecko), choosing them based on closely related species in OHNOLOGS v2 database (see Materials and Methods for details). Across the three species, approximately 20-30% of vertebrate-specific (Vertebrata phylostratum) hub genes are retained as ohnolog pairs (Table 1). This pattern is most pronounced in zebrafish and lungfish, congruent with previous results showing that sauropsids have highest lineage-specific loss of 2R ohnologs (*10*). By comparison, genome-wide analyses estimate that ohnologs constitute on average 25% of the extant gene set of vertebrate genomes (*10*), in line with the general trend for the vertebrate genomes. This indicates that while hubs are not particularly overrepresented among retained ohnologs, their retention echoes the overall influence of WGD and dosage sensitivity on the evolution of vertebrate gene networks.

Natural selection is one of the primary evolutionary forces shaping evolutionary transitions, filtering genetic variation in response to new environmental pressures and favoring traits that enhance survival and reproductive success (*43*, *44*). Among the different forms of selection, directional selection plays a central role during habitat shifts, driving the fixation of alleles that confer functional advantages under novel conditions (*45*).

In order to investigate whether directional selection shaped the evolution of hub genes key for lungfish estivation, we first tested which HOGs had undergone significant shifts in directional selection in two evolutionary episodes: the branch leading to vertebrates, and the one leading to Sarcopterygii (including lungfish and tetrapods). Next, we checked which of those HOGs contained at least one hub gene of lungfish under estivation. We identified 91 directionally selected hub HOGs in the branch leading to Vertebrates and 18 in the branch leading to Sarcopterygii lineages.

Among the 91 directionally selected hub-containing HOGs in the vertebrate branch that orchestrate lungfish estivation, several fell into intermediate KEGG categories linked to adaptive structural and physiological change (Fig. 4a). Some of these included proteins involved in membrane trafficking (e.g., Annexin A6, Arf-GAP domain protein 1) (*46*, *47*); proteolysis and protein turnover (e.g., calpains, serpin), supporting cytoskeletal reorganization and tissue protection (*48*, *49*); autophagy and stress response (e.g., Sestrin 1/3), a key regulator of metabolic suppression and oxidative stress defense (*50*) and cytoskeleton and motility regulation (e.g., transgelin), essential for vascular tone and organ structural and tissue remodeling (*51*). Together, these categories highlight selective pressures on cellular systems that enhance respiratory, circulatory, and metabolic adaptability, traits likely advantageous during the water-to-land transition.

**Fig. 4.**
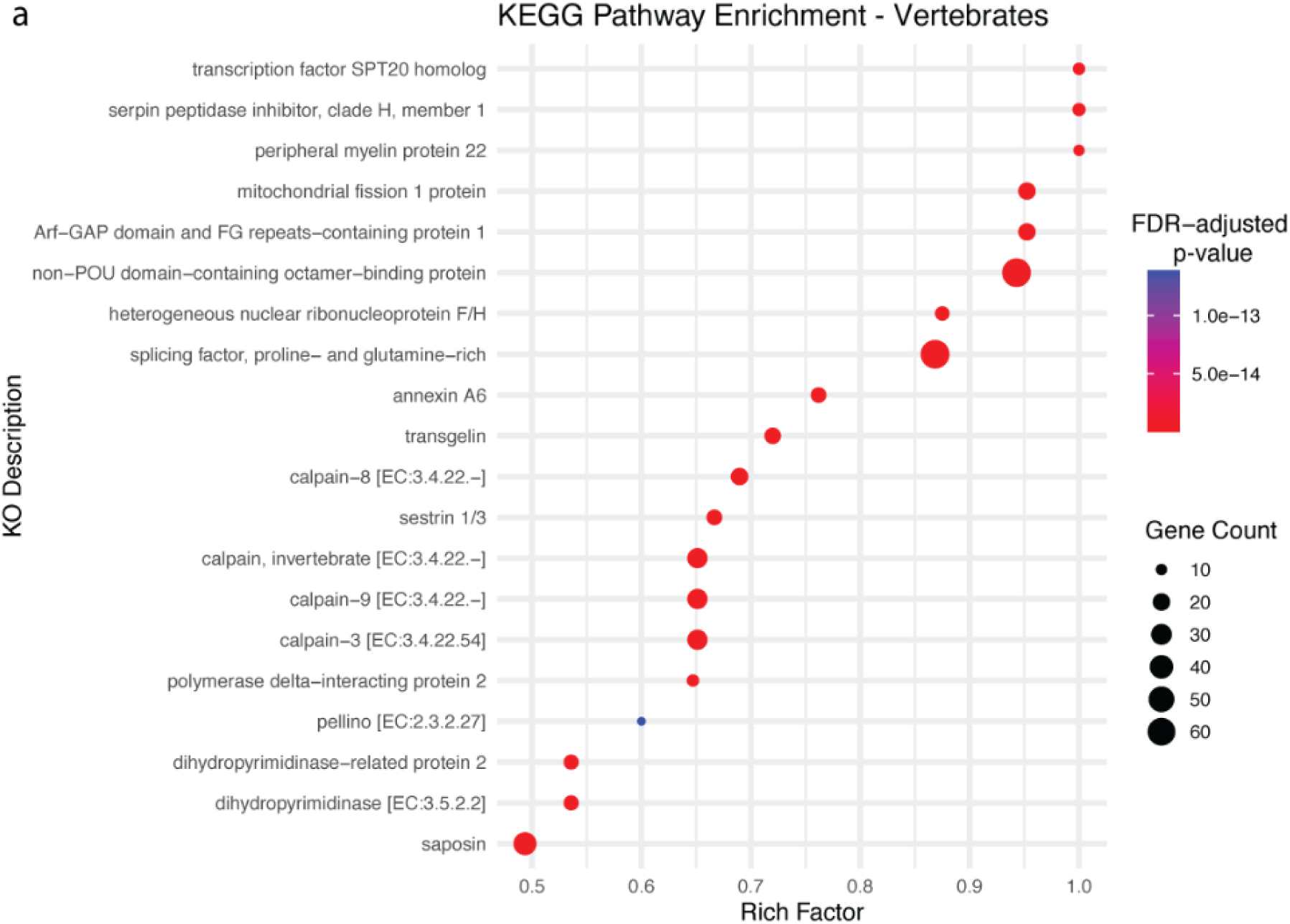
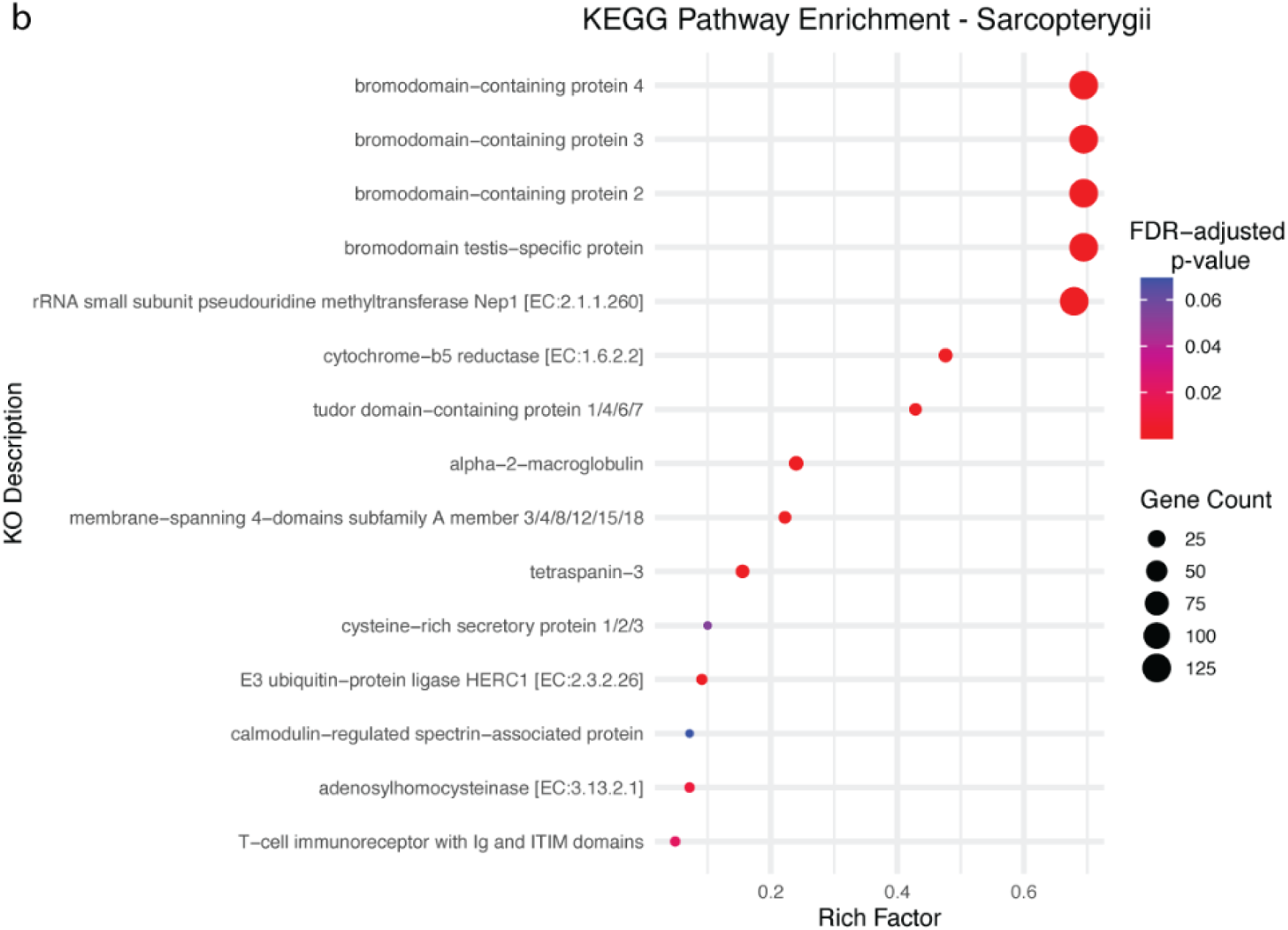
KEGG pathway enrichment of directionally selected hub-containing HOGs in internal vertebrate phylogenetic branches. Bubble plots show significantly enriched KEGG pathways for hub-containing HOGs under directional selection in the (a) branch leading to Vertebrata, and (b) branch leading to Sarcopterygii. The y-axis lists the KEGG Orthology (KO) term descriptions, and the x-axis indicates the Rich Factor (i.e. ratio of hub-containing HOGs annotated to a given pathway related to the total number of genes in that pathway). Bubble size reflects the number of genes associated with each KO term, and bubble color indicates the false discovery rate (FDR)–adjusted *p*-value, with red representing more significant enrichment.

Among the 18 directionally selected hub HOGs in the Sarcopterygii branch, several mapped to KEGG categories relevant to cellular and regulatory processes (Fig. 4b). Some of these included membrane trafficking and cell adhesion (e.g., Tetraspanin-3) (*52*); ubiquitin-mediated proteolysis and signaling (e.g., E3 ubiquitin–protein ligase HERC1) (*53*); and epigenetic regulation and transcriptional control (e.g., bromodomain-containing proteins) (*54*). Interestingly, some of these pathways also play critical roles in immunity such as bromodomain-containing proteins which are master transcriptional regulators of innate immune genes (*55*). Together, these categories indicate selective pressures on cellular and regulatory systems that may have supported physiological plasticity and environmental responsiveness in the lineage leading to tetrapods.

We next asked whether ohnologs important for estivation also exhibit signatures of directional selection. To address this, we intersected (i) the set of hub genes identified before as retained ohnologs correlated with lungfish under estivation and (ii) the directionally selected HOGs included hub genes correlated with estivation in lungfish assigned also to the Vertebrata phylostratum. Our analysis revealed a group of retained ohnologs that occupy hub positions in gene co-expression networks and are simultaneously subject to directional selection, totalling 21 such HOGs. This pattern points to a plausible evolutionary scenario in which ancient ohnologs, preserved through WGD, have been co-opted, or exapted, to facilitate some of the physiological demands of estivation.

The 21 identified hub ohnologs relevant for estivation of lungfish are enriched for functional categories (Fig. 5) associated with membrane trafficking and structural maintenance (e.g., Annexin A6, p24 family protein alpha) (*46*, *56*), water transport and osmotic regulation (e.g., Aquaporin-1, Aquaporin-4) (*57*), cytoskeletal regulation via Rho signaling (e.g., Rho GTPase-activating proteins 12/27, 15, 9; microtubule-associated protein RP/EB family) (*58*), stress response and metabolic regulation (e.g., Sestrin 1/3, Cyclophilin B) (*50*, *59*), and RNA processing (e.g., hnRNP F/H) (*60*). Additional categories include ubiquitin-mediated signaling (e.g., RNF130) (*61*), mitochondrial dynamics that have been found to be activated in response to stress induced by hypoxia (e.g., FUN14 domain–containing protein 1) (*62*), and phosphoregulation (e.g., phosphohistidine phosphatase) (*63*). The prominence of these categories is consistent with previous work showing that estivation in African lungfish skin relies on early pro-inflammatory and antimicrobial responses, and suggests that similar immune-linked programs may operate system-wide during dormancy (*64*). Several of the identified hub ohnologs, including Annexin A6, aquaporins, Cyclophilin B, Rho GTPase-activating proteins, sestrins, RNF130, and hnRNPs, have established roles in immune cell activation, cytokine signaling, and regulation of inflammatory pathways, indicating that estivation recruits ancient networks at the interface of stress physiology and immunity. Notably, key immune effectors previously described in the lungfish cocoon, such as antimicrobial peptides, pro-inflammatory cytokines, mucins, and granulocyte markers, are embedded within significant estivating modules, directly linking our hub-centric networks to the inflammatory barrier and tissue-protective responses characterized in skin (*64*). In this context, the combined involvement of these genes in membrane trafficking, extracellular matrix dynamics, and cytoskeletal remodeling points to a coordinated module that not only preserves water balance and cellular homeostasis but also supports controlled inflammation, tissue repair, and regeneration during repeated water–land transitions.

**Fig. 5.**
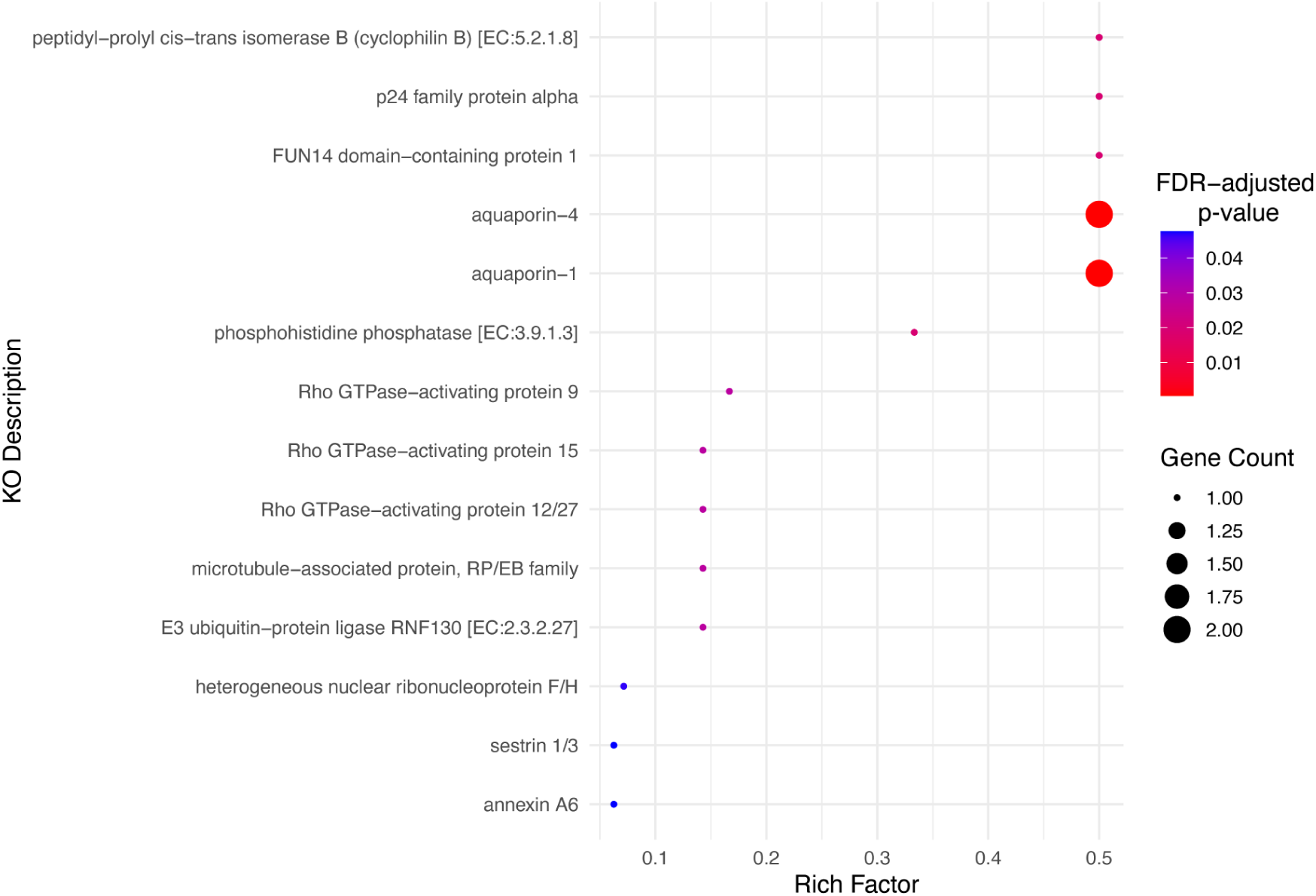
KEGG pathway enrichment of hub ohnologs key for estivation in lungfish showing shifts in directional selection in Vertebrata phylostratum. Bubble plots showing significantly enriched KEGG pathways for hub-containing HOGs under directional selection that are also retained ohnologs of the 2R WGD in the vertebrate branch. The y-axis lists the KEGG Orthology (KO) term descriptions, and the x-axis indicates the Rich Factor (ratio of hub-containing HOGs annotated to a given pathway to the total number of genes in that pathway). Bubble size reflects the number of genes associated with each KO term, and bubble color indicates the false discovery rate (FDR)–adjusted *p*-value, with red representing more significant enrichment.

Previous work in African lungfish skin revealed the essential role for pro-inflammatory immune responses early on during estivation (*64*). Beyond the skin, several hub ohnologs identified here indicate a potential role for immunity in whole body adaptation to terrestrial life in African lungfish. For instance, Annexin A6 plays diverse roles in inflammation and cytokine secretion (*65*) and cyclophilin B attracts diverse immune cells participating in tissue inflammation (*66*). Additionally, Rho GTPase activating proteins are critical molecular switches for immune cell migration, cytokine production and antimicrobial immune responses (*67*). Moreover, sestrins can resolve chronic inflammation by suppressing pro-inflammatory cytokine responses via autophagic pathways (*68*). As inflammation is inherently linked to tissue remodeling and repair (*69–71*), our findings may indicate organ-wide co-option of hub ohnologs that support physiological inflammatory responses required for tissue remodeling during water-land transitions. Together, these functions suggest that retained ohnologs under directional selection have been co-opted into coordinated systems for water conservation, tissue remodeling, inflammatory responses and cellular stress resilience during estivation in lungfish. The intersection of the 2R retained ohnologs, hub genes, and directionally selected genes highlights how WGD set the stage for later evolutionary innovation. Our findings could possibly support a model of exaptation, where the legacy of WGD provided the raw material for adaptive refinement under new environmental pressures.

Taken together, our results illustrate how major genomic events like WGD can have long-term evolutionary consequences by equipping lineages with genetic reservoirs that are subsequently shaped by selection to produce novel and extreme phenotypes under challenging ecological conditions, such as the out-of-the-sea transition in vertebrates.

Our study shows that the molecular foundations of vertebrate terrestrialization are rooted in deeply conserved gene networks that predate the water-to-land transition, but which later were shaped by duplication and selection. By integrating multi-tissue transcriptomic data from lungfish, including estivating states, into a comparative co-expression framework across vertebrates, we suggest a three-step hypothesis underlying the evolution of the gene repertoire potentially facilitating vertebrate terrestrial adaptation: first, ancestral co-expression modules provided a core regulatory toolkit; second, whole-genome duplications supplied additional dosage-sensitive hubs that were retained; and third, directional selection subsequently refined these hubs to meet the physiological challenges of life on land. In lungfish, these functions converge in the physiology of estivation, suggesting that traits originally serving survival under drought may have been co-opted as preadaptations for terrestrial life. Together, these findings support a model in which the water-to-land transition was not driven by de *novo* genetic innovation alone, but by the co-option and directional refinement of ancient, duplication-enriched networks involved in membrane trafficking, water balance, cytoskeletal regulation, and stress response under new ecological pressures. More broadly, this study illustrates how ancient genome-scale innovations can be repeatedly repurposed by selection to generate the complex physiological phenotypes that enable vertebrates to colonize and thrive on land.

## Materials and Methods

### Sample collection

Different tissues for European sea bass, zebrafish, lungfish, natterjack toad and the moorish gecko were dissected (see Table S1). For lungfish, tissues from specimens under estivation were also sampled. To complement and strengthen our dataset we also used a publicly available dataset of lungfish gills and lungs of control and estivating samples (PRJNA903405). All African lungfish experiments and tissue collection were performed under the University of New Mexico IACUC protocol #25-201710-MC. Experimental estivation was performed as previously described by us (*18*, *64*). Of note, while animals were bled from the caudal vein prior to tissue sampling, not all blood was removed and therefore we cannot rule out the presence of blood in several tissues including the brain and gills. For European sea bass, fish were euthanised with an overdose of MS222 and death confirmed by severing the spinal cord prior to tissue dissections. Zebrafish and lungfish were sacrificed using cold thermal and spinal cord transection. For the natterjack toad (*Epidalea calamita*), euthanasia was carried out by immersing the toads for 1 h in a 5 g/L solution of MS222 buffered with pharmaceutical-grade sodium bicarbonate to pH 7.5 at room temperature (*72*).

Death was confirmed by severing the spinal cord prior to tissue dissections. For the Moorish gecko (*Tarentola mauritanica*), euthanasia was performed by intramuscular administration of alfaxalone (Alfaxan, 10 mg/kg) to induce deep anesthesia, followed by spinal cord severing to ensure death prior to dissection. All amphibian and reptiles specimens used in the study were collected and processed under permit SF/0206/23 issued by the Generalitat de Catalunya, Departament d’Acció Climàtica, Alimentació i Agenda Rural, Direcció General de Polítiques Ambientals i Medi Natural. All procedures complied with Spanish regulations (RD53/2013) and European Directive 2010/63/EU for animal experimentation, and were approved by the Local Government of the Generalitat de Catalunya (ref. 10209)

### RNA Extraction, Sequencing and Preprocessing

RNA extractions were performed using the TRIzol® reagent (Invitrogen, USA) method following the manufacturer’s instructions and using MaXtract® High Density tubes (Qiagen) to help with solvent removal and minimize contamination following mechanical sample homogenization either by plastic rotor pestles or cryogenical disruption using ceramic mortar and pestle, depending on the sample. The concentration of all samples was assessed by Qubit RNA BR Assay kit (Thermo Fisher Scientific). Libraries were prepared with the TruSeq Stranded mRNA library preparation kit (Illumina), and sequenced on a NovaSeq 6000 (Illumina, 2 × 150 bp) for a minimum of 6Gb coverage. Summary statistics of sequenced data are shown in Tables S1-S5.

Although a subset of species in our dataset have well-characterized, high-quality genomes, others do not. To enable robust and comparable analyses across all species, we generated full-length transcriptomes using Iso-Seq, which have been shown to sufficiently capture gene repertoire evolutionary dynamics, particularly when completeness metrics such as BUSCO scores are high (*73*). For each species, the same RNA extractions used for short-read sequencing were equimolarly pooled to prepare the Iso-Seq libraries used as reference datasets in co-expression network analysis.

RNA samples were subjected to DNAse treatment using the Turbo DNA-free DNase (Invitrogen) following the manufacturer’s instructions. SMRTbell libraries were generated following the procedure ‘Preparing Iso-Seq® libraries using SMRTbell® prep kit 3.0 (PN 102-396-000 REV02 APR2022)’. To enable pooling of multiple samples on 1 SMRTcell, libraries were made using barcoded adapters. The libraries were sequenced on a Sequel-IIe using Sequel II sequencing kit 2.0 and Binding kit 3.1 with 24 hr movie-time. Summary statistics of sequenced data is shown in Table S7.

Raw Illumina RNA-seq reads for all the tissues and species mentioned above were preprocessed using fastp (-q 10 -u 50 -y -g -Y 10 -e 20 -l 100 -b 150 -B 150) to remove adapters and low quality reads. Trimmed reads were assessed using FastQC (https://www.bioinformatics.babraham.ac.uk/projects/fastqc) before any further analysis. For Iso-Seq data processing, reads were demultiplexed using the IsoSeq pipeline v4.0.0 (https://github.com/PacificBiosciences/pbbioconda), and cDNA primers were removed with LIMA v2.7.1. PolyA tails and artificial concatemers were trimmed using isoseq refine (v4.0.0). Full-Length Non-Concatemer (FLNC) reads were then clustered into de novo reference transcriptomes using CD-HIT v4.8.1 (cd-hit-est, a=0.99) (*74*). Each species-specific high-quality reference transcriptome was evaluated using BUSCO v5.4.7 with the metazoa_odb10 database (*75*), and all achieved high BUSCO completeness scores (Table S7), supporting their suitability for downstream analyses.

Transcriptomes were translated into protein sequences using TransDecoder v5.5.0 (https://github.com/sghignone/TransDecoder), and decontaminated with BlobTools v2 (https://blobtoolkit.genomehubs.org/blobtools2/). The longest protein isoform was extracted using a custom script and retained for subsequent functional analyses. These longest isoform proteomes were then used as input for orthology inference (see below).

### Gene co-expression network construction

To build gene co-expression networks, trimmed Illumina RNA-seq reads from each tissue were quasi-mapped to their respective species-specific Iso-Seq reference transcriptomes using Salmon v1.10.2 (*76*). Genes with at least 10 counts in 90% or more of samples were retained, minimizing data sparsity. Expression data were then transformed using the regularized log (rlog) method from DESeq2 v1.44.0 (*77*), which stabilizes variance across a broad range of expression values and is well suited for co-expression network inference.

To control for unwanted technical variation, batch effects were identified and corrected using the removeBatchEffect() function in limma v3.60.6 (*78*), incorporating surrogate variable analysis to account for hidden batch structure. Data quality was assessed by removing low-quality genes and outlier samples with the goodSamplesGenes() function from WGCNA v1.73 (*26*). Additional diagnostic checks - including principal component analysis and sample clustering via heatmaps - were conducted using plot_PCA() and plot_heatmaps() functions from BioNERO v1.12.0 (*79*), prior to downstream analysis.

Gene co-expression networks were independently generated for each species using Weighted Gene Co-expression Network Analysis (WGCNA), based specifically on RNA-seq data derived from each tissue. The soft-thresholding power (β) was selected using the pickSoftThreshold() function, applying biweight midcorrelation (bicor) and a signed hybrid network type to approximate scale-free topology (R² ≥ 0.8). Network topology and connectivity were validated using softConnectivity() and scaleFreePlot() functions.

Final networks were constructed with the blockwiseModules() function (TOMType = signed, networkType = signed hybrid, deepSplit = 4, minModuleSize = 30, mergeCutHeight = 0.15). Module color assignments were extracted for further analysis. Modules showing significant correlation with the tissues (p < 0.05) were retained, and hub genes were identified as those with high gene significance (|GS| > 0.2) and high intramodular connectivity (|KME| > 0.8).

### Orthology inference

Hierarchical orthologous groups (HOGs) were inferred using FastOMA v0.4.4 (*28*) using the omamer database for LUCA and default parameters. Longest isoform proteins from sea bass, zebrafish, lungfish, natterjack toad and common wall gecko were used as the primary species, and *Ciona intestinalis* (Urochordata), *Branchiostoma belcheri* (Cephalochordata), *Lytechinus variegatus* (Echinodermata), *Saccoglossus kowalevskii* (Hemichordata) serving as outgroups. Outgroups’ longest isoform proteins were downloaded from MATEdb2 (*80*). The evolutionary age of each HOG (i.e. positioning them in the internal nodes of the phylogenetic tree) was estimated with pyHam (*81*).

### Phylostratigraphic reconstruction of hub genes

To reconstruct the evolutionary history of hub genes, we mapped the species-specific hub transcripts from each condition to their corresponding proteins in the reference proteomes and assigned them to HOGs for phylostratigraphic analysis using custom scripts (see ‘Code availability’). Since each species had a different number of hub genes and these linked to varying numbers of HOGs, we normalised the data by calculating percentages, i.e. dividing the number of genes assigned to each phylostratum (Node) by the total number of hub genes per species. This approach ensured comparability across species and prevented bias due to differences in gene set sizes.

### Jaccard similarity analysis

To evaluate the conservation of co-expression modules across species, we assessed the overlap between the gene contents of lungfish statistically significant modules in regard to tissues under estivation and those of the other species (sea bass, zebrafish, natterjack toad and common wall gecko) using the Jaccard similarity index. This analysis was based on shared HOGs, allowing comparisons across species irrespective of gene identifiers. After assigning the genes to HOGs for each module in lungfish, we computed the Jaccard similarity index with every module from the other four species. The resulting pairwise Jaccard scores thus represent the degree of shared evolutionary content between modules across species at the HOG level. Since there is no standard threshold for defining module conservation based on Jaccard similarity in this context, we adopted a data-driven threshold. Specifically, we selected a cutoff of > 0.07, which corresponded to the highest range of similarity values observed in our dataset, allowing us to focus on the most strongly overlapping module pairs. These modules were then subjected to downstream functional analysis to determine whether shared HOG content reflected conservation of biological functions across vertebrates.

### Ohnolog identification

To assess whether the observed burst of hub-containing HOGs at the base of the vertebrate lineage could be explained by the retention of ohnologs from 2R-WGD, we performed an ohnolog enrichment analysis focused on that branch. We selected three representative vertebrate species from our dataset that span aquatic, transitional, and terrestrial lifestyles and had a closely related species in OHNOLOGS v2 database (*10*): zebrafish, lungfish and common wall gecko. For each species, we identified putative ohnologs. We used reciprocal BLASTp searches with MMseqs2 (*82*) against a corresponding reference proteome included in OHNOLOGS v2 database: *Da. rerio* against *Da. rerio* reference proteome, *P. annectens* against *L. oculatus*, and *T. mauritanica* against *A. carolinensis.* Ohnologs were defined as reciprocal best hits retained in both species pairs. For each species we downloaded ohnologs with intermediate q-score.

We then mapped the identified ohnologs to their corresponding HOGs and examined the overlap with hub-containing HOGs (i.e., those including hub genes from statistically significant modules for estivation detected in lungfish co-expression network). We focused specifically on HOGs that originated at the Vertebrata phylostratum.

After identifying the intersection between hub-containing HOGs and retained ohnologs from vertebrate WGDs, we compared this set with the intersection of HOGs under directional selection (inferred as described below) to highlight gene families shaped by both duplication and adaptive evolutionary pressures.

### Directional selection analysis

To investigate selective pressures associated with key evolutionary transitions in vertebrates, particularly those involving shifts in habitat along the aquatic–terrestrial transition, we employed Pelican v1.0.8 (*83*). We focused on two internal nodes of the species phylogeny (Vertebrata and Sarcopterygii), which represent successive divergence events predating the emergence of tetrapods and their transition to terrestrial environments. For each node, we selected HOGs that originated at the branch leading to it and retained at least one gene from each of the descendant clades and always including lungfish, ensuring a balanced taxonomic representation for comparative analysis and a maximum representation of our focal species.

Each selected HOG was first quality-filtered using PREQUAL v1.02 (*84*) to remove unreliable sites, followed by multiple sequence alignment with MAFFT v7.525 (*85*). Alignments were then trimmed with trimAl v1.5 (*86*) to retain only the most informative regions. Since the alignment and trimming steps occasionally led to loss of key sequences, we re-applied the original taxonomic criteria to ensure that each HOG still contained at least one representative from every descendant clade defined at the node. HOGs that no longer met this requirement were excluded from further analysis.

The final set of filtered HOGs was used to infer gene trees with IQ-TREE2 v2.4.0 (*87*) using the LG4M model. These gene trees were then reconciled with the species tree using Treerecs v1.2 (*88*) to account for duplication and loss events. The reconciled trees, along with their corresponding alignments, were input into Pelican to test for signatures of directional selection changes across amino acid sequences. For each HOG, we defined the terrestrial-adapted species *E. calamita* and *T. mauritanica* as the foreground, and the remaining species as the background character. This allowed us to assess whether lineage-specific selective pressures were associated with adaptation to terrestrial environments. Using Gene-wise Truncated Fisher’s method (*89*) in the output of Pelican, which is a set of p-values for each site in every alignment, we aggregated site-level p-values into gene-level predictions by considering the first k=10 p-values per alignment. We further applied the (*90*)

At each node, we then examined how many of the HOGs under directional selection also contained hub genes identified in lungfish. This allowed us to assess whether genes showing evidence of directional selection were involved in key regulatory roles within the co-expression networks of estivating lungfish. Specifically, we quantified the overlap between HOGs under selection and those containing at least one hub gene from any lungfish estivating tissue.

### Functional annotation and pathway enrichment analysis

Functional annotation of selected HOGs from different subset datasets was performed with eggNOG-mapper v2.1.10 (*91*) using the eggNOG DB v.5.0.2 (*92*). KEGG pathway assignments from the eggNOG-mapper output were then used for enrichment analysis with (*93*). For each subset enrichment test, lungfish genes were used as the background, with the background set adjusted to the most appropriate gene set each time (Table S19).

## Author contributions

Conceptualization: KE, RF

Sample provision: RH, IS, SC, LR, MB

Methodology: KE, JSO, NE, CVC, RF

Investigation: KE

Visualization: KE

Supervision: RF

Writing - original draft: KE, RF

Writing - review & editing: KE, JSO, NE, CVC, RH, LR, MB, SC, IS, RF

## Data and materials availability

Data, code and scripts are available at the Github repository https://github.com/MetazoaPhylogenomicsLab/Eleftheriadi_et_al_2025_Lungfish_comparative_genomics/ *(to be released before publication).* Transcriptomic data is publicly available in SRA under BioProject PRJNA1025375 (*to be released before publication*).

## Supplementary material

Supplementary Figures can be found here: [Supplementary Material.pdf](https://drive.google.com/file/d/1tly4BzxDBJDDdgjk97IvGEjWOwKSHh9x/view?usp=share_link) and supplementary tables can be found here: [Supplementary_Tables](https://docs.google.com/spreadsheets/d/13tQ2xxH36HrQvF5Yxl82gSY5iezVJNF7BPjkEsNzoZI/edit?gid=727658808#gid=727658808).

